# Cardiac synchrony during collaborative drawing: A longitudinal comparison of same generation and intergenerational dyads

**DOI:** 10.1101/2025.10.23.684101

**Authors:** Ryssa Moffat, Luca A. Naudszus, Emily S. Cross

**Affiliations:** Social Brain Sciences Lab, ETH Zurich

**Author notes:** Correspondence: Ryssa Moffat, Emily S. Cross.

**Keywords:** intergenerational, collaboration, drawing, physiological synchrony, relationship formation

## Abstract

Intergenerational social programs provide opportunities for people of all ages to form new relationships. Furthermore, existing qualitative and behavioural evidence from such programs points to health and wellbeing benefits, yet the physiological consequences of repeated intergenerational encounters remain unknown. A deeper understanding of how such programs shape dyadic physiological responses will illuminate the mechanisms of relationship formation. Across a six-session collaborative drawing program, we tracked cardiac synchrony within 31 intergenerational (older/younger adult) and 30 same generation (younger adult) dyads. Each session dyads completed self-report measures, then drew together and alone, while we recorded participants’ actions with motion capture and physiological signals (neural and cardiac) using fNIRS. Collaborative behaviour, self-reported social closeness, and interpersonal distance (i.e., proximity) showed group-specific patterns, whereby interpersonal distance emerged as a promising objective measure of relationship development. Cardiac synchrony did not covary with group, task, an interaction thereof or any measure of behaviour or social closeness–yet there was a trending relationship between collaboration while drawing together and cardiac synchrony for intergenerational dyads only. In summary, cardiac synchrony pointed to marginally enhanced arousal during active collaboration between older and younger adults. Relationship development was better characterised, in this study, by behaviour and self-report measures than cardiac synchrony.

## 1 Introduction

In response to worrying reports of declining social connection, many communities have begun to employ intergenerational arts programs to improve community members’ health and wellbeing ^[1,2]^. Growing behavioural and qualitative data from such programs suggests that intergenerational arts programs offer much needed opportunities to build social connection, reduce stereotypes about other generations, and provide structures for people to engage in activities that add meaning to their lives ^[2–6]^. Intergenerational arts programs that offer repeated contact older and younger generations, as opposed to one-off sessions, seem to yield the greatest improvement in social connectedness and feelings of affiliation between generations ^[5]^. More broadly, feelings of social connectedness or closeness (e.g., being strangers vs. friends vs. romantic partners) are known to modulate physiological signals (e.g., cardiac signals and brain activity) during collaboration ^[7]^. Yet, longitudinal evidence tracking the extent to which physiological signals index relationship formation is only just beginning to emerge ^[8]^. A deeper understanding of how repeated intergenerational collaboration fosters relationship formation, across experiential and physiological levels, holds potential to inform program structure and resource allocation for intergenerational arts programs. Moreover, new insights into intergenerational collaborative behaviour can only strengthen efforts to promote healthy ageing.

In this study, intergenerational and same generation dyads completed a 6-week collaborative drawing program, in which they met as strangers and interacted with the same partner for all 6 sessions. We examined how collaboration and physiological synchrony, specifically *cardiac synchrony*, evolved across sessions. Specifically, we tracked how cardiac synchrony levels during collaboration were shaped by repeated social encounters, the generational constellation (intergenerational vs. same generation), as well as dyads’ feelings of social closeness and interpersonal distance (i.e., physical proximity to one another).

### 1.1 Cardiac synchrony an index of shared arousal during collaboration

The cardiovascular system is reactive to changes in both psychological and physiological factors ^[9]^. Fluctuations in heart rate reflect changes in arousal of the central nervous system resulting from changes in an individual’s environment ^[10]^. Increases in heart rate reflect sympathetic activity, which is associated with excitement and stress. Reductions in heart rate reflect parasympathetic activity, relating to relaxation and divided attention ^[11,12]^. When people gather in groups of two or more, changes in their physiological signals including heart rate reflect a mixture of self-regulation and co-regulation ^[13,14]^, which can result in greater temporal alignment of heart rates, per se, and heart rate variability among two or more people. Precise time-locking of fluctuations can arise from external environmental factors–take for example a jumpscare (i.e., a startling event) in a horror movie causing a spike in cinemagoers’ heart rates simultaneously. Time-lagged alignment occurs more commonly in live social interaction involving behavioural coordination, for example turn taking in conversations ^[15]^. Such alignment in arousal can result from shared attention ^[16]^, emotional contagion ^[17,18]^, or joint action ^[19,20]^. In other words, measures of temporal alignment of heart rates and heart rate variability between individuals, or cardiac synchrony, can shed light on shared fluctuations in physiological arousal during social interactions. Accordingly, shared fluctuations in arousal, tracked using measures of cardiac synchrony, are reported to underpin important aspects of collaboration, including perceptions of group cohesion, satisfaction, commitment, subjective togetherness, trust, and perceived empathy (see review by Mayo and colleagues ^[7]^).

Several studies have investigated the extent to which cardiac synchrony can predict the outcomes of collaboration ^[19,21–24]^. Sharika et al. ^[24]^ recorded 44 groups of 204 students completing a verbal consensus-seeking task. The authors reported that cardiac synchrony between group members predicted the likelihood of reaching consensus significantly better than individuals’ heart rates, self-reported ratings of team function, or duration of the consensus-seeking process. On this basis, Sharika et al. proposed that cardiac synchrony may be useful as a biomarker of engagement during collaboration ^[24]^. Gordon et al. ^[22]^ compared levels of cardiac synchrony during synchronised drumming, freely improvised drumming, and resting state (i.e., no movement). They reported that cardiac synchrony was greater during drumming than baseline but did not differ between drumming conditions, highlighting the role of joint action in aligning fluctuations in arousal. Gordon et al. further reported that levels of cardiac synchrony during synchronised drumming predicted levels of coordination of drumming in the subsequent improvised drumming task. Alignment of arousals levels, as indexed by cardiac synchrony, may thus be a precursor for successful collaboration ^[22]^.

Other studies suggest that perceptual shifts during collaboration drive changes in shared arousal ^[19,21,23]^. Noy et al. tracked levels of perceived and kinematically measured togetherness when dyads played a mirror game. Controlling for differences in movement intensity, the authors found that cardiac synchrony increased during periods of togetherness, more so for perceived than kinematic togetherness. Periods of togetherness were also notably accompanied by increased heart rates, reflecting increased arousal ^[19]^. Boukarras et al. ^[21]^ recorded grasp timing from dyads told to grasp two bottle shaped objects according to specific instructions. In half the trials, the instructions prompted the same or complementary grasping behaviours, and in the other half of trials, the instructions prompted evoked peer-to-peer or leader-follower grasp dynamics. None of these prompted grasping behaviours showed a statistically significant relationship between grasping timing and cardiac synchrony. However, Boukarras et al. did observe heightened levels of cardiac synchrony when dyads shifted between peer-to-peer to a leader-follower dynamics or from same to complementary movements. These findings led the authors to suggest that perceptual shifts during collaboration dynamics, rather than specific patterns of behavioural coordination, may influence arousal levels, and by extension, cardiac synchrony ^[21]^. Findings from a study by Mitkidis et al. ^[23]^ lend support to the idea that perceptual shifts during collaboration may alter shared arousal. Mitkidis et al. compared cardiac synchrony in a group that completed interleaved collaborative Lego building tasks with public goods games to a group that completed only Lego building tasks. They report that the public goods game led to greater cardiac synchrony during Lego building. They also found that cardiac synchrony predicted the participants’ expectations of their partners during the public goods game. Mitkidis et al. interpret both results as being reflective of increased awareness of the partner relating to trust-building processes. Considered together, these findings highlight that perceptual shifts during collaboration impact arousal levels as quantified by cardiac synchrony. Notably, too much shared arousal can have deleterious effects on collaboration ^[25,26]^, suggesting that co-regulation of arousal levels is central to successful collaboration ^[7]^.

Changes in familiarity that occur within or across encounters may represent a sufficient perceptual shift to influence shared arousal, and by extension, cardiac synchrony. Fusaroli et al. ^[27]^ invited five groups of 4-5 students to build Lego models in 5-minute blocks, alternating between building alone and together as a group, and computed measures of heart rate coordination (i.e., the similarity and stability of signals within groups–complementary measures to cardiac synchrony). The authors reported that heart rate coordination within groups increased from the first block of collaborative model-building to the last and that coordination of movement and speech within groups predicted heart rate coordination ^[27]^. These findings illustrate how increasing familiarity, on a short time scale, co-occurs with increasing alignment of behaviour and arousal levels. Highlighting the relevance of understanding changes in dyadic levels of shared arousal over repeated encounters, Gernert et al. ^[28]^ measured cardiac synchrony during psychotherapy sessions targeting affective disorders at two sessions, approximately two weeks apart. The authors found a positive association between cardiac synchrony across both sessions and changes in self-reported depressive symptom severity between the two sessions ^[28]^. Unfortunately, this study does not report the difference in cardiac synchrony between sessions, leaving the trajectory of shared arousal an open question. While empirical attention has been paid to first encounters ^[e.g., 29]^, little is known about the trajectory of fluctuations in shared arousal across repeated encounters. To advance our understanding of the prerequisites for relationships formed over repeated encounters to blossom into robust social bonds and the conditions that predict relationships fizzling out, we need to document the trajectories of shared arousal across multiple encounters alongside measures of relationship quality. By considering these trajectories in same generation and intergenerational dyads, we can provide insights for optimising relationship formation in intergenerational community programs.

### 1.2 Relationship quality constrains shared arousal, and by extension, cardiac synchrony

Relationship quality and shared fluctuations in arousal, as measured by cardiac synchrony, are known to covary, though the effect size derived from a meta-analysis of 26 studies is quite small ^[7]^, and the directionality of the effect is not known. Many of the studies conducted have been dedicated to understanding shared arousal in romantic partners and are summarised in Mayo et al.’s meta-analysis ^[7]^. For example, watching emotional videos together has been shown to elicit greater cardiac synchrony between strangers, relative to friends and romantic partners ^[17]^. Bizzego et al. propose that greater familiarity between friends and romantic couples results in their autonomic systems being less attuned to changes in the other person’s autonomic responses. This may be a feature of the task involving non-interactive copresence with both people’s attention directed to the videos. Other tasks, involving co-presence and interaction show the opposite pattern. At a firewalking performance, Konvalinka et al. ^[30]^ recorded cardiac synchrony between firewalkers and their friends and between firewalkers and strangers. The authors compared these groups and reported greater cardiac synchrony between firewalkers and friends and family members than strangers. The authors similarly suggest that social closeness ^[30]^ is a driving factor in the coupling of autonomic responses. Focusing on first-time encounters, Adel et al. ^[29]^ recorded cardiac synchrony in dyads comprising strangers who shared emotional stories. Dyads showed greater cardiac synchrony when they self-reported high levels of mutual liking. Relationship quality seems to have a more nuanced impact on shared arousal when collaboration is compared with competition. Danyluck and Page-Gould ^[11]^ demonstrated that self-reported perceived similarity and interest in friendship between dyad members can shape cardiac synchrony levels differentially for collaboration and competition. Whereas cardiac synchrony is positively associated with perceived similarity in both conditions, cardiac synchrony is positively associated with interest in friendship only during collaboration and not during competition ^[11]^. Recently, Andersen et al. ^[31]^ recorded cardiac synchrony between visitors who went through a recreational commercial haunted house in groups, designed to induce fear in the visitors. The authors reported that cardiac synchrony is closely related to individuals’ ratings of subjective arousal and that cardiac synchrony was greater between dyads within each visitor group who were socially close than those who were not. The findings contribute to the evidence that social relationships shape co-regulation through shared fluctuations in arousal ^[31]^. Based on these studies, we would expect that in interactive contexts evoking average physiological arousal (i.e., non-frightening; not involving haunted houses or risky behaviour such as firewalking), cardiac synchrony would be highest at a first encounter and decrease as dyads who began as strangers become acquainted and socially closer.

Self-reported feelings of social closeness to another person tends to result in individuals being comfortable in closer physical proximity to each other ^[32–34]^. Like social closeness, physical proximity (henceforth interpersonal distance) is bidirectionally related with shared arousal ^[31,35]^. In a 2×2 design, Jackson et al. ^[35]^ recorded groups of people walking freely vs. in synchrony as a group vs. manipulated arousal levels by having the leader walk above (high arousal) or below (low arousal) average walking speed. The researchers found that the pack was the tightest (least distance between walkers) during the condition combining synchronous and high arousal walking. The researchers propose that their findings reflect how arousal and synchrony can be employed to manufacture group cohesion ^[35]^. To our knowledge, no studies to date investigate the relationship between shared arousal measured via cardiac synchrony and interpersonal distance. Nonetheless, previous research using other measures of physiological synchrony has revealed that arousal levels are likely associated with interpersonal distance. For example, teams of paramedic trainees who show greater synchrony in electrodermal activity work physically closer to one another and engage in more cooperative dialogue compared to teams showing lower synchrony ^[36]^. Interpersonal distance appears to be associated with shared arousal levels during collaboration, yet additional evidence is needed to establish this relationship’s reliability.

### 1.3 Cardiac synchrony in intergenerational relationships is poorly documented

Notably, the relationships between shared arousal, social closeness, interpersonal distance and collaboration have (to our knowledge) only been examined in dyads consisting of two young adults at a single point in time (as opposed to longitudinally). In very few cases, intergenerational patterns of shared arousal have been explored within mother-child dyads (see review by DePasquale and colleagues ^[37]^). Older adults have yet to be represented in such studies, a trend that is observed in experimental studies more broadly ^[38,39]^. It is critical to remedy the underrepresentation of older adults as the proportion of older adults in the world’s population is steadily climbing ^[40]^ and older adults are at elevated risk of experiencing loneliness ^[2]^. Moreover, longitudinal insights extending beyond first encounters and covering stages before friendships and romantic relationships are necessary for a deeper understanding of the physiological underpinnings of relationship development.

### 1.4 Present study

This research sought to shed light on the physiological mechanisms of collaboration in budding relationships–both same generation and intergenerational, including older adults. We preregistered (https://osf.io/xuvy5) the three following experimental aims:

- Document the trajectory of cardiac synchrony over the course of six sessions of collaborative drawing within both same generation and intergenerational dyads.
- Explore how levels of collaboration visible in drawings (evaluated by external raters) relate to dyad members’ feelings of social closeness, physical closeness, and performance on a separate collaborative task (i.e., completing a jigsaw puzzle).
- Explore how cardiac synchrony is related to levels of collaboration visible in drawings. Specifically, we wish to assess whether this relationship differs between same generation and intergenerational dyads, and the extent to which this relationship is impacted by dyad’s feelings of social closeness, physical closeness, and performance on a separate collaborative measure (i.e., jointly completing a jigsaw puzzle).

In reporting these findings, charting cardiac synchrony within intergenerational and same generation dyads across 6 sessions in an everyday context similar to a community art program, we reveal first insights into longitudinal patterns of shared arousal, in a unique design that involves older adults. The implications of this work are central to understanding the behavioural and physiological mechanisms governing relationship formation and collaboration across the lifespan. Moreover, this work sheds light on important considerations regarding the implementation of cardiac synchrony as an objective measure of social connection or collaboration in everyday, real-world settings.

## 2 Methods

### 2.1 Participants

#### Main sample

We recruited 122 community-dwelling participants from Zurich, Switzerland (31 older adults, aged 69+ years; 91 younger adults, aged 18–35 years). Included participants were fluent in German and had no known history of neurological or psychological disorders (e.g., stroke, concussion, ADHD, autism, schizophrenia, or depression). Participants were assigned to intergenerational dyads (*n* = 31) or same generation dyads (*n* = 30) based on availability for sessions (e.g., matching people available at the same day/time for 6 consecutive weeks). All dyads started the study as strangers. Intergenerational dyads comprised one older adult (76 ± 4 years); 18 female, 13 male) and one younger adult (24 ± 4 years; 19 female, 11 male, 1 other). Same generation dyads comprised two younger adults (22.5 ± 3.6 years; 36 female, 24 male). Supplementary Table 1 details the number of individual participants’ recordings and dyads’ sessions that were excluded, as well as reasons for exclusion.

Our recruitment campaign involved advertising through the University of Zurich’s Healthy Longevity Centre, ETH Zurich’s DeSciL student pool, clubs and organisations for senior citizens (e.g., theatre groups and choirs/orchestras), and social media. All participants provided written informed consent prior to participation. Ethical approval was obtained from the Ethics Committee of the Canton Zurich (Ref: 2023-01073). The study was conducted according to the Declaration of Helsinki. We report how we determined our sample size, all data exclusions, all manipulations, and all measures in the study.

#### External ratings of drawings

To obtain ratings of drawings produced by the main sample (described further in 2.3.1), we recruited 148 new participants via Prolific after the main study concluded. To be included, participants had to be over18 years old, proficient English speakers, residents of the United Kingdom. To ensure response quality, we only accepted participants who had previously completed a minimum of 50 tasks on Prolific with a 100% approval rate. Of these participants, 45 were excluded for failing to respond to all attention checks correctly (i.e., by moving all sliders all the way to the right of the sliding scale when prompted). We thus use ratings from 103 raters in our analyses. We did not collect demographic information from these participants. All participants provided written informed consent. Ethical approval was obtained from the Ethics Committee of ETH Zurich (Ref: 2024-N-265). The study was conducted according to the Declaration of Helsinki.

### 2.2 Procedure

Each dyad completed six weekly sessions (once per week, for six weeks). At each session, participants completed questionnaires querying feelings of loneliness ^[41]^ and attitudes toward their own and the other (older or younger) generation ^[42]^. Participants also completed a questionnaire measuring empathy ^[43]^ at the first session. Dyads drew alone once and collaboratively twice with oil pastels provided by Caran D’Ache, for a duration of five minutes (Figure 1; Supplementary Figure 1). Participants were not given any prompts regarding what to draw or how to draw (e.g., no suggestions for motifs). Participants were instructed not to talk while drawing, so that speech would not be a confound in comparisons of drawing alone (of particular relevance for neural and cardiac measures). Participants were instructed that they could talk freely during the rest of the session. After drawing, participants completed a collaborative activity, with different activities each week. In the third session, the collaborative activity was to complete a 54-piece jigsaw puzzle. Activities completed on other weeks include sorting pastels, a divergent thinking task, and a joint verbal fluency task. While participants drew and completed collaborative activities, we recorded their cortical activity using functional near-infrared spectroscopy (fNIRS) over the bilateral inferior frontal gyri (IFG) and bilateral temporo-parietal junction (TPJ). Furthermore, we video-recorded participants’ movement using a GoPro camera to capture movements of the head and hands. At the end of each session, after we stopped the fNIRS and video recordings, participants completed the Inclusion of Other in Self scale ^[44]^. Analyses of fNIRS recordings, loneliness scores, and attitudes toward generations are reported elsewhere, as are additional specifics regarding the fNIRS recording methodology ^[8]^.

**Figure 1.**
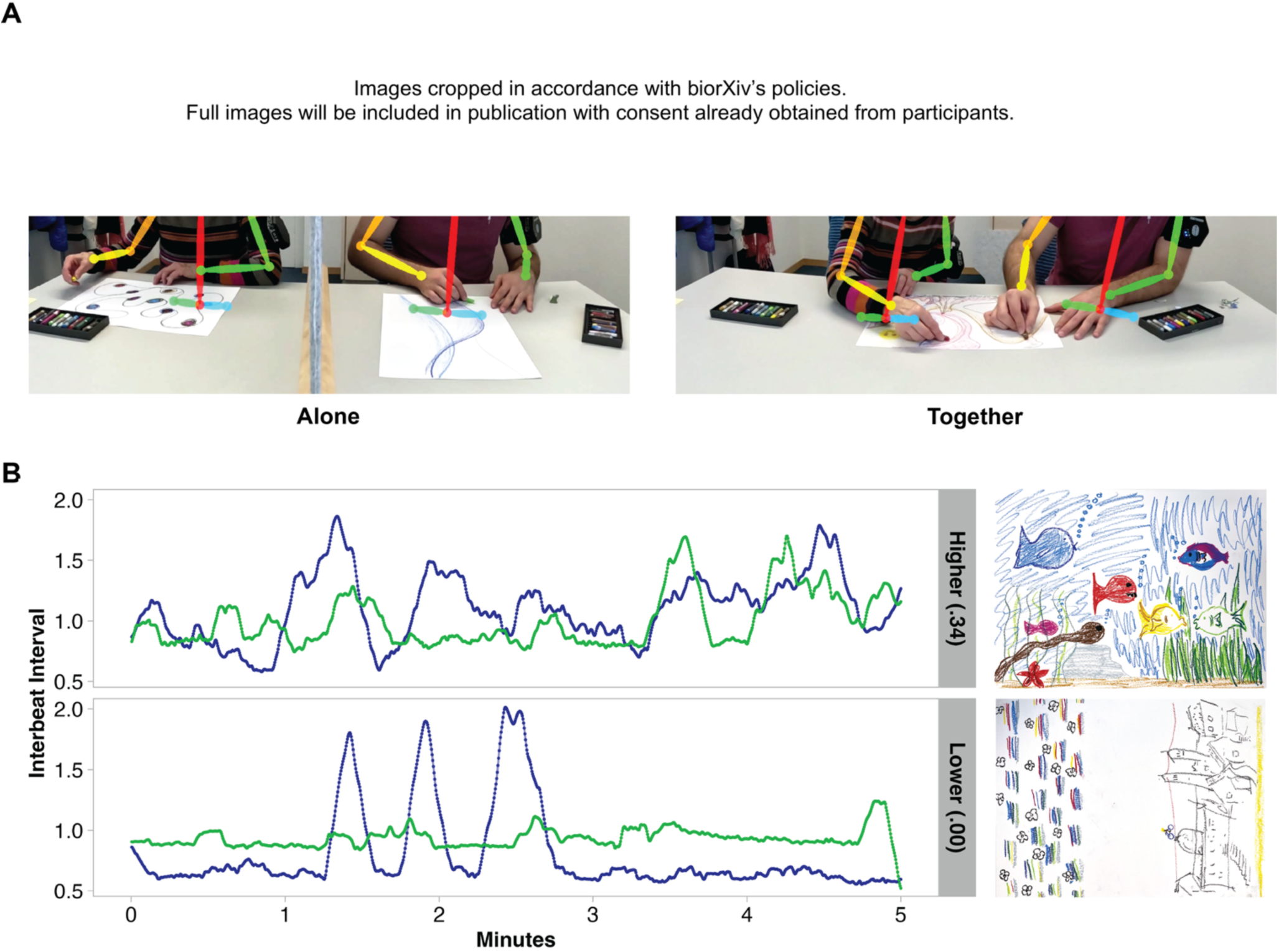
A) Left: An intergenerational dyad drawing independently on separate pieces of paper during fNIRS recording. The dyad members are separated by a grey felt divider that obscures their view of each other. Right: The same intergenerational dyad drawing together on a single piece of paper during fNIRS recording. The colourful lines overlaid on top of the participants show the 2D motion tracking used to capture interpersonal distance, specifically the distance between dyad members’ neck joints. B) IBI times series of two different dyads, illustrating high and low levels cardiac synchrony with the produced drawing (see further examples sorted by collaboration ratings in Figure 2).

### 2.3 Measures

#### 2.3.1 Drawing collaboration

Each drawing that the dyads created was judged by independent raters recruited via Prolific. Raters (*n* = 103) were instructed that they would view 122 drawings, each created by two people at once on a single piece of paper. They were asked to rate the degree of harmony between the elements of the drawings (on a sliding scale from 0 “entirely independent” to 100 “entirely coordinated”) and the use of space (on a sliding scale from 0 “divided” to 100 “unified”). To obtain a measure of the perceived coherence of the drawing, which we propose reflects the level of collaboration during drawing, we summed the ratings of harmony and use of space. Thus, raw scores of drawing collaboration could range from 0-200. Examples of drawings with highest, average, and lowest collaboration ratings are shown in Figure 2.

**Figure 2.**
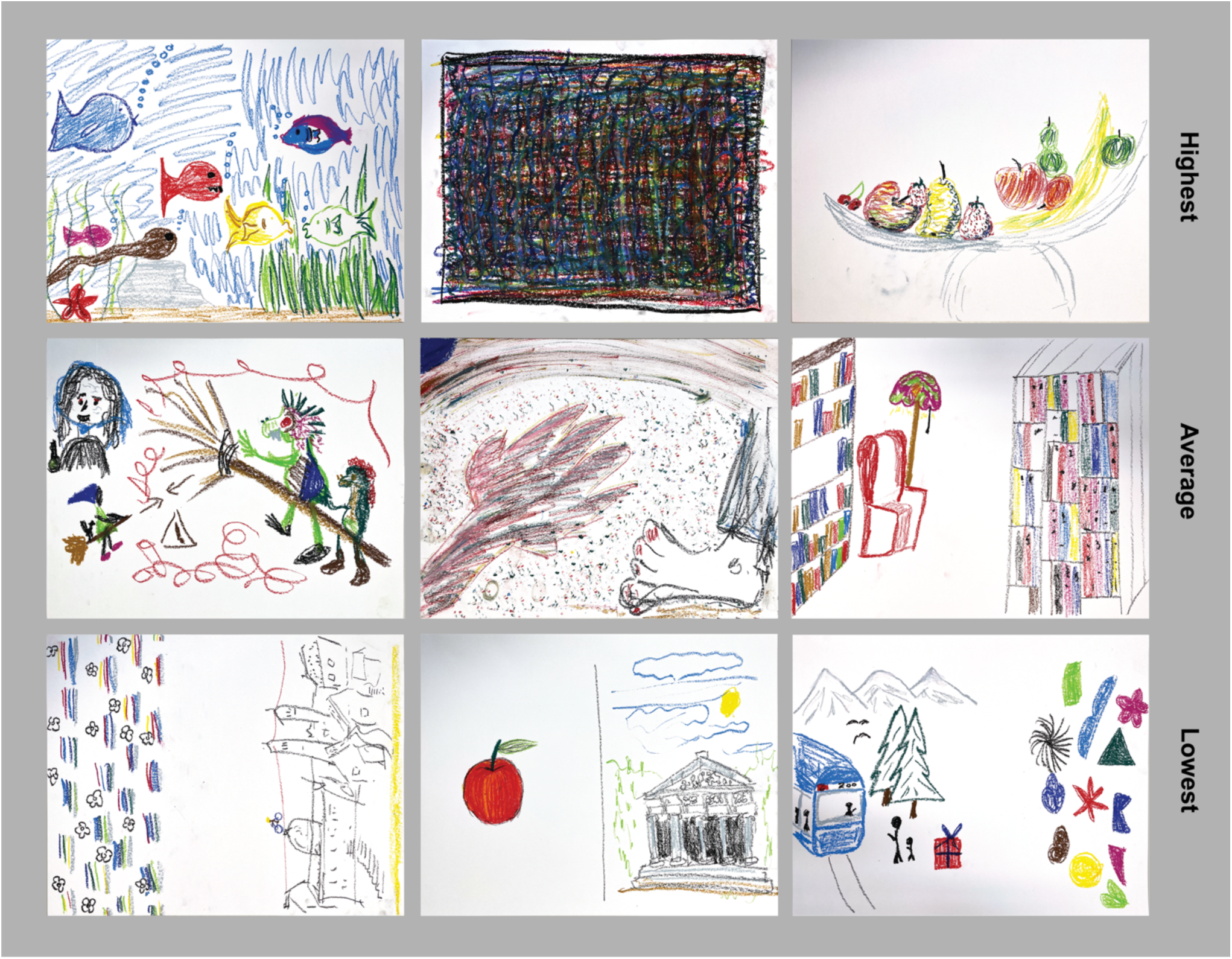
Examples of drawings organised by rating of collaboration. Top row = highest collaboration ratings; middle row = average collaboration ratings; bottom row = lowest collaboration ratings. Ratings are derived by summing raters’ ratings of the harmony between the elements of the drawings (0 “entirely independent” to 100 “entirely coordinated”) and use of space (0 “divided” to 100 “unified”). The left-most drawings in the top and bottom row are also presented in Figure 1B alongside the IBI time series recorded while the drawings were made.

#### 2.3.2 Puzzle collaboration

After completing the drawing component of the third session, dyads completed a puzzle as the collaborative activity for that session. Speaking was permitted. Dyads had six minutes to complete a 54-piece jigsaw puzzle of cartoon animals in jungle (pieces’ dimensions: ∼ 8 × 8 cm). Dyads’ scores corresponded to the number of pieces assembled in 6 minutes. In the case that participants assembled all pieces in less than 6 minutes, the remaining number of seconds to 6 minutes was divided by 6.66 (the average number of seconds per piece if completed in 6 minutes; 360 s /54 piece = 6.66 s) and added to a score of 54.

#### 2.3.3 Social Closeness

To understand how collaborative behaviour shapes feelings of social closeness, we administered the Inclusion of Other in Self (IOS) scale (Aron et al., 1992 after dyads drew together and completed the final collaborative task. Participants viewed seven sets of overlapping circles representing the self and the partner, ranging from no overlap to near complete overlap of the two circles, and indicate which set of circles best represents their relationship with their partner (i.e., the other dyad member). To capture cumulative social closeness within dyads and differences in feelings of social closeness within dyads, we computed summed and absolute difference IOS scores per dyad ^[21,45]^. We refer to these measures as social closeness (Sum) and social closeness (Dif).

#### 2.3.4 Interpersonal distance

We used the videos recorded at each session to track upper body movements while dyads drew together. Specifically, we used OpenPose ^[46]^ to estimate 2D pose per frame, returning coordinates for predifined body parts. To assess interpersonal distance (sometimes referred to as proximity in other work), we computed the average distance between the coordinates of each dyad members’ neck joint for each 5-minute instance of drawing together. In our analyses, this measure is referred to as distance.

### 2.4 Data analysis

#### 2.4.1 Extracting inter-beat intervals

As per our preregistration, we extracted the cardiac signal by identifying peaks in HbO concentrations ^[47]^. We cleaned the HbO signal from neural and further physiological components such as Mayer waves by upsampling it to 100 Hz and applying a zero-phase third-order Butterworth IIR band-pass filter ^[48]^ with cut-offs at 0.5 and 2.5 Hz ^[49]^. Then, we smoothed the signals using a Savitzky-Golay filter with window length 15 and a 5^th^-order polynomial. We identified peaks with the AMPD algorithm ^[47]^, implemented in the python *ampdlib* package ^[50]^, and transformed the signal peak timestamps to an inter-beat interval (IBI) time series, smoothly interpolated at 4 Hz. We originally preregistered ARIMA cleaning ^[51]^, but in practice it reduced the signal to near-zero fluctuations, largely removing the slow structure that may reflect the physiology of interest in this context. Given the lower signal-to-noise ratio of fNIRS-derived IBIs when compared to ECG (electrocardiography) signals, and our total number of samples (fNIRS signals recorded at ∼ 5 Hz), we instead analysed IBI series on their original scale to retain physiologically meaningful variance.

#### 2.4.2 Calculating synchrony

We calculated cross-correlation functions (CCFs) between all dyads’ cleaned physiological signals in each session and activity across time lags of +/- 3 seconds (−12 to +12 lags at 4 Hz) and retained non-absolute CCF values. Examples of time series yielding high and low cardiac synchrony estimates are visualised in Figure 1B.

#### 2.4.3 Group-level analysis

We completed our preregistered exploratory statistical analyses using a Bayesian approach to multilevel regression ^[52]^. We fit models using the brms package ^[version 22.2;, 53]^ in the R language ^[version 4.5.1;, 54]^ within the RStudio IDE ^[version 2025.05.1+513;, 55]^. We report and interpret the posterior distribution of relevant model parameters using a 95% credible interval, calculated via the 95% highest posterior density region (HPD) method ^[52]^. For readers familiar with Frequentist approaches that include p-values, we recommend perusing Kruschke & Liddell ^[56]^, and we share a (simplified) heuristic for interpreting HPDs: Parameter and comparison estimates can be considered *substantial* when the HPD does not contain zero and can be considered trends when the tip of an HPD-tail overlaps with zero.

##### Measures of collaboration, social closeness, and interpersonal distance

To assess the extent to which collaboration, social closeness, and interpersonal distance changed across the six sessions, as well as relationships between measures, we fit a series of models in the following structure: *MeasureA ∼ 1 + MeasureB * Group * Session + (1|DyadID)*. We extracted point estimates for all sessions combined and slopes across sessions, and subsequently computed contrasts between groups (intergenerational vs. same generation).

##### Cardiac synchrony for all sessions combined and across sessions

We first sought to establish if cardiac synchrony levels differed substantially between real and pseudo dyads. We fit the following model: *CardiacSynchrony ∼ DyadType * Group * Task * Session*. We did not include random intercepts at this stage, as the total number of real and pseudo dyads exceeded 6000. Random intercepts, accounting for dyad-level variance, were included in subsequent group level analyses.

Contrasts between real and pseudo dyads were computed to confirm that some component of the cardiac synchrony was attributable to real social interaction, as opposed to similar experiences or homeostatic biological rhythms.

##### Relationships between cardiac synchrony and measures of collaboration, social closeness, and interpersonal distance

First, we fit a model to the cardiac synchrony data, which we averaged across all lags. The model was: *CardiacSynchrony ∼ Group * Drawing Collaboration + Group * Puzzle Collaboration + Group * Social Closeness (Sum)+ Group * Social Closeness (Dif) + Group * Interpersonal Distance + Session + (1|DyadID)*. Subsequently, we fit a model to examine relationships between cardiac synchrony at separate lags and our measures. Models were structured as follows and fit per measure: *CardiacSynchrony ∼ Lag * Group * Measure + (1|DyadID)*. For both steps, we extracted point estimates for all sessions combined and slopes across sessions, and subsequently computed contrasts between groups (intergenerational vs. same generation).

## 3 Results

### 3.1 Behavioural and self-report measures

#### 3.1.1 Contrasts between groups

See Table 1 for estimates and 95% HPD (highest posterior density region) for each measure. Analyses of drawing collaboration, puzzle collaboration and interpersonal distance are original to this study. Social closeness (Sum) and (Dif) have previously been analysed by our research team ^[8]^; we include these analyses here for completeness. **Drawing collaboration.** Drawing collaboration differed substantially between groups, with the intergenerational drawings showing greater collaboration than same generation drawings. Drawing collaboration did not change substantially across sessions for either group, though there was a trend of intergenerational dyads showing a greater increase in collaboration than same generation dyads. **Puzzle collaboration.** Puzzle collaboration differed markedly between groups, with same generation dyads scoring substantially higher than intergenerational dyads. **Social closeness (Sum).** Summed social closeness was substantially greater for intergenerational dyads than same generation dyads. Summed social closeness increased substantially across sessions for both groups at a similar rate. **Social closeness (Dif).** Intergenerational dyads showed substantially greater differences in dyadic social closeness than same generation dyads. Dyadic differences in social closeness decreased across sessions substantially for same generation dyads and showed the same trend for intergenerational dyads.

**Table 1.** Standardised estimates computed per behavioural and self-report measure with HPD in square brackets. Point estimates for all six sessions combined are presented first, followed by the estimates of the slope of change across sessions in the lower section. No slope of change is available for puzzle collaboration, as dyads only completed the puzzle task once, at the third session.

|  | Drawing<br>Collaboration | Puzzle<br>Collaboration | Social<br>Closeness (Sum) | Social<br>Closeness (Dif) | Interpersonal<br>Distance |
| --- | --- | --- | --- | --- | --- |
| **Sessions combined** |  |  |  |  |  |
| Intergenerational | 0.20 [0.08, 0.32] | -0.50 [-0.59, -0.42] | 0.17 [0.7, 0.26] | 0.13 [0.03, 0.23] | 0.07 [-0.18, 0.03] |
| Same gen | -0.03 [-0.15, 0.08] | 0.45 [0.36, 0.53] | -0.18 [-0.27, -0.08] | -0.08 [-0.18, 0.03] | -0.06 [-0.04, 0.17] |
| Intergen-Same gen | 0.23 [0.07, 0.40] | -0.95 [-1.07, -0.84] | 0.35 [0.22, 0.47] | 0.20 [0.07, 0.34] | -0.13 [-0.28, 0.00] |
| **Change across six sessions** |  |  |  |  |  |
| Intergenerational | 0.04 [-0.01, 0.93] | - | 0.22 [0.17, 0.27] | -0.05 [-0.10, 0.00] | 0.02 [-0.02, 0.07] |
| Same gen | -0.01 [-0.07, 0.04] | - | 0.16 [0.11, 0.21] | -0.07 [-0.13, -0.02] | 0.07, [0.01, 0.12] |
| Intergen-Same gen | 0.05 [0.00, 0.11] | - | 0.09 [-0.02, 0.20] | 0.02 [-0.03, 0.07] | -0.04 [-0.10, 0.00] |

##### Interpersonal distance

Interpersonal distance showed a trending group difference, with same generation dyads showing substantially lower interpersonal distance than intergenerational dyads. Interpersonal distance increased substantially across sessions for same generation dyads only. Comparisons between groups showed a trending group difference, where the slope of change in interpersonal distance across sessions was more positive for same generation than intergenerational dyads.

#### 3.1.2 Relationships between measures

Relationships between measures of collaboration, self-reported social closeness, and interpersonal distance per group are illustrated in Figure 3. Drawing collaboration was substantially positively associated with puzzle collaboration for the intergenerational group only, yielding a substantial difference between intergenerational and same generation dyads (ß = 0.26, [0.08, 0.42]). Drawing collaboration was substantially positively associated with summed social closeness for both groups, but the association was stronger for same generation dyads, relative to intergenerational dyads (ß = - 0.16, HPD = [-0.30, -0.02]). Drawing collaboration was not substantially associated with differences in dyadic social closeness. Drawing collaboration was substantially negatively associated with interpersonal distance for intergenerational dyads and substantially positively associated with interpersonal distance for same generation dyads, yielding a substantial group difference (ß = -0.21, HPD = [-0.25, -0.07]). Puzzle collaboration was substantially negatively associated with summed social closeness for same generation dyads only, resulting in a substantial group difference (ß = 0.18, HPD = [0.06, 0.30]). Puzzle collaboration was also substantially negatively associated with differences in dyadic social closeness for same generation dyads only, resulting in a substantial group difference (ß = 0.21, HPD = [0.09, 0.33]). Puzzle collaboration was substantially negatively associated with interpersonal distance for same generation dyads only, and no substantial group difference was observed. Summed social closeness was substantially positively associated with dyadic differences in social closeness for both groups and did not differ substantially between groups. Interpersonal distance was substantially positively associated with summed social closeness for intergenerational dyads and substantially negatively associated with summed social closeness for same generation dyads, with a substantial difference between groups (ß = 0.22, HPD = [0.08, 0.36]). Interpersonal distance showed a trending negative association with differences in dyadic social closeness for intergenerational dyads and a substantial positive association for same generation dyads, yielding a substantial difference between groups (ß = -0.20, HPD = [-0.35, 0.06]).

**Figure 3.**
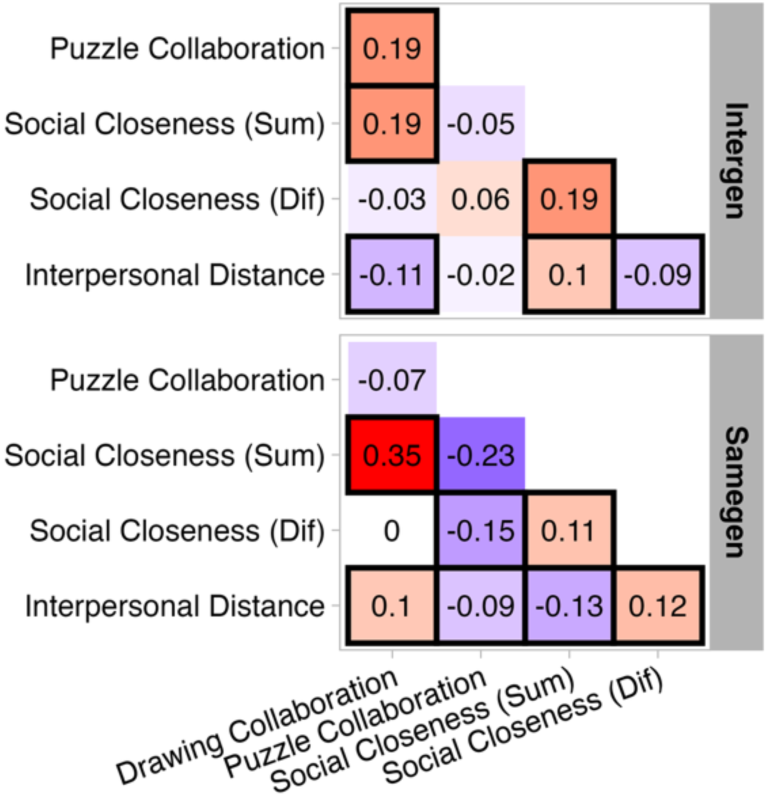
Standardised estimates of relationship between behavioural and self-report measure per group. Black squares indicate that relationships are substantial (i.e., 95% HPD does not overlap with zero). Note: the relationship between Interpersonal Distance and Social Closeness (Dif) for same generation dyads is trending, rather than substantial, but in a black square to facilitate interpretation of results. Standardised parameter estimates per measure shown with 95% HPD in Supplementary Table 2.

In plain language, these findings can be summarised as follows: Intergenerational dyads’ collaboration was similar across different tasks. Dyads who drew more collaboratively also assembled more pieces of the jigsaw puzzle collaboratively. For both groups, dyads’ summed social closeness predicted their level of drawing collaboration (i.e., both members feeling very close co-occurred with higher drawing collaboration). Intergenerational dyads who sat physically closer together made more collaborative-looking drawings, whereas the same was true for same-generation dyads who sat further apart. Intergenerational dyads who sat closer together reported feeling socially closer to each other, whereas same-generation dyads who sat further apart reported feeling socially closer to each other.

### 3.2 Cardiac synchrony

#### 3.2.1 Comparing real and pseudo dyads

**All sessions combined.** For all lags averaged and separate lags, we observed no substantial difference in cardiac synchrony levels between real and pseudo dyads for drawing together or alone (Supplementary Figure 2). **Across sessions.** For all lags averaged and separate lags, the rate of change in cardiac synchrony across sessions (all lags averaged) did not differ substantially between real and pseudo dyads for drawing together or alone (Figure 4; Supplementary Figure 3).

**Figure 4.**
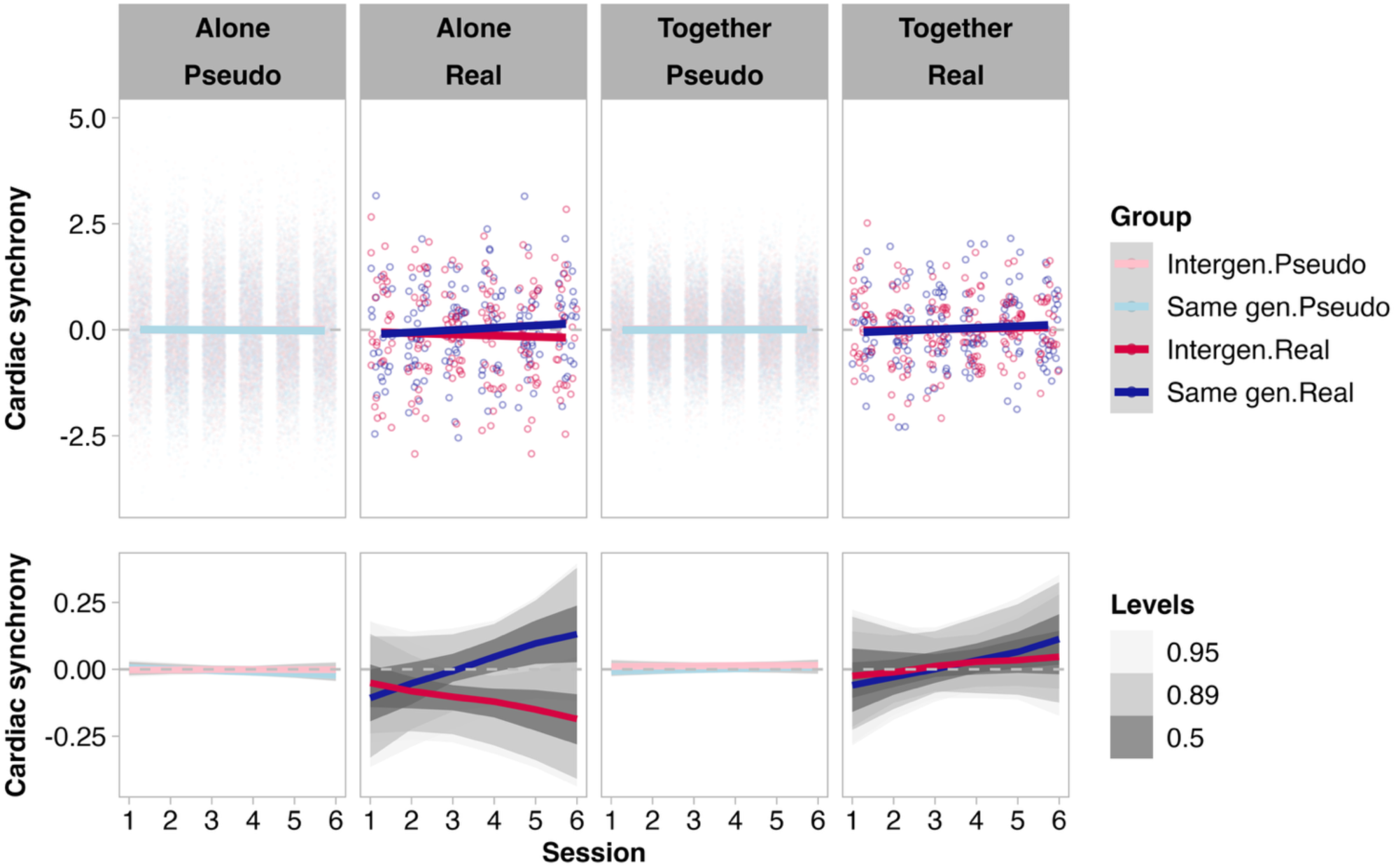
Trajectories of cardiac synchrony in real and pseudo dyads across sessions. Top panel shows individual data points (dots) and estimated slopes (lines). Bottom panel zooms in on the estimated slopes, with grey shading showing 95%, 89%, and 50% intervals of the posterior predictive distributions. Neither the overall level of cardiac synchrony for all sessions combined nor the rate of change in cardiac synchrony across sessions differed substantially between real and pseudo dyads. Though real dyads’ cardiac synchrony the trajectories across sessions may appear to differ, the 95 % HPD overlaps > 10 % (threshold for a trend, Section 2.4.3) for drawing alone and together.

We interpret these findings as follows: Any two individuals meeting the inclusion criteria, who experience the specific dyadic experimental environment applied here would show comparable levels of cardiac synchrony. True interaction with an assigned partner does not appear to contribute uniquely to cardiac synchrony levels. We present the further analyses in accordance with our preregistration.

#### 3.2.2 Comparisons between drawing conditions and groups

**All sessions combined.** For all lags averaged, we observed no substantial effects of drawing task, group or task*group interactions on cardiac synchrony levels. **Across sessions.** For all lags averaged, we observed no substantial main effect (i.e., no substantial change in cardiac synchrony levels across sessions) and no substantial effects of drawing task, group or task*group interactions on the slope of cardiac synchrony across sessions.

### 3.3 Relationships between cardiac synchrony, behaviour, and self-report measures

As per our preregistration, we first explored relationships between cardiac synchrony with all lags averaged, collaborative behaviour, social closeness, and distance. We found no substantial relationships between cardiac synchrony and either measure of collaboration, social closeness or interpersonal distance. Also in line with our preregistration, we explored relationships between cardiac synchrony and each measure per lag (n = 25; from -3000 ms to 3000 ms in 250 ms increments). We observed no substantial relationships for any measure and no substantial difference between groups at any lag (Figure 5). For drawing collaboration, intergenerational dyads showed a trending positive relationship between cardiac synchrony and drawing collaboration at all lags except -3000 ms, -2750 ms, -2500 ms, and -1750 ms. Note: 95% HPD with no overlap with 0 indicates a slope substantially different than 0, < 10 % overlap with 0 indicates a trending difference from 0. At the four lags that did not show trending differences, the degree of 95 % HPD overlap with 0 ranged from 10-14 %. For intergenerational dyads, positive lags reflect the older dyad member’s IBI signal leading or coming earlier in time, and negative lags reflect the younger member’s IBI signal leading or coming earlier in time. As we observed trending positive and negative lags trending, there is no evidence that a dyad member of a certain generations fills a leader role.

**Figure 5.**
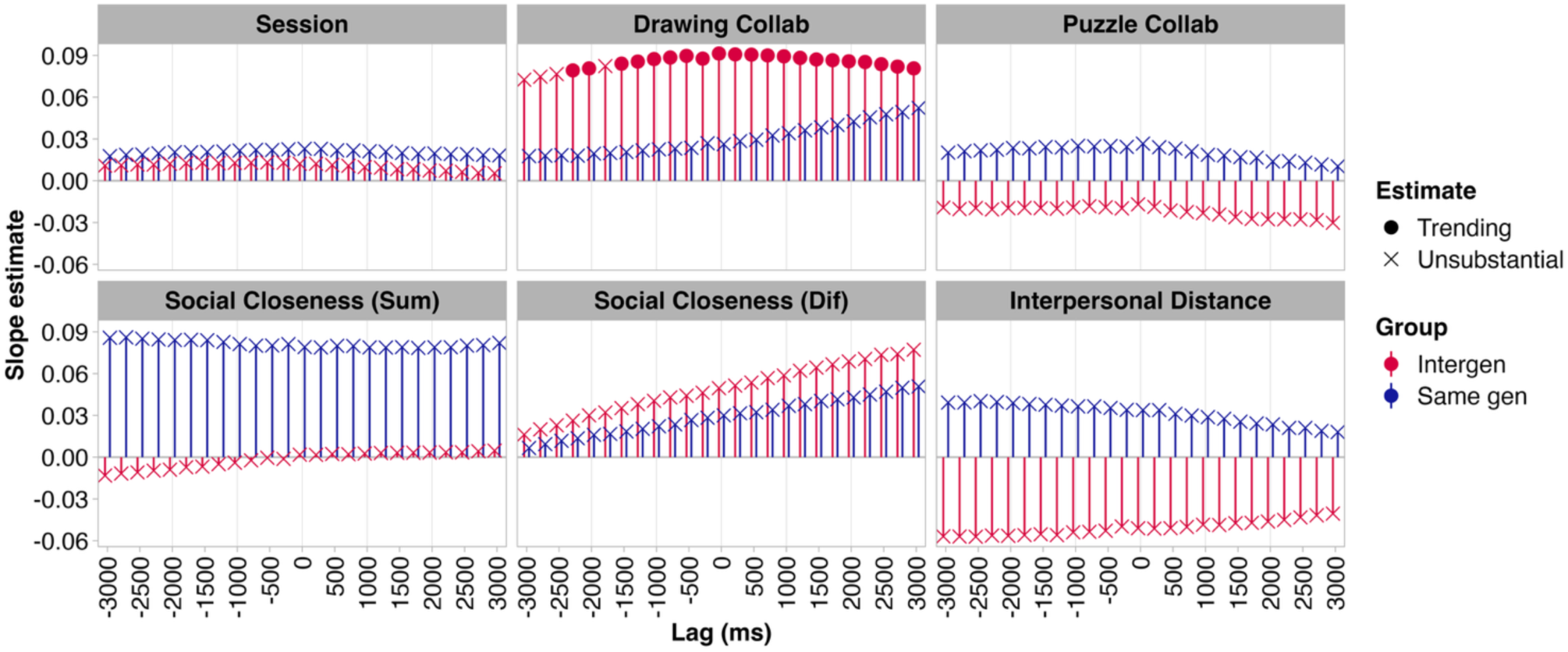
Standardised estimates of relationship between cardiac synchrony and session number, as well as behavioural measures of collaboration, self-reported measures of social closeness, and interpersonal distance per lag (n = 25; from - 3000 ms to 3000 ms in 250 ms increments). No substantial relationships observed. Drawing collaboration panel shows trending relationships between cardiac synchrony and drawing collaboration scores for intergenerational dyads at all but 4 lags. See Supplementary Figure 4 for visualisation including error bars showing 95 % HDP.

## 4 Discussion

In this study, we charted cardiac synchrony, collaborative behaviours and social closeness in intergenerational and same generation dyads across 6 sessions in an everyday context resembling a community art program. Dyads’ collaborative behaviours and measures of social closeness showed nuanced relationships, primarily characterised by greater collaboration in cases of greater social closeness. For cardiac synchrony–both averaged and separate lags–we found no substantial differences between real and pseudo dyads overall or across sessions, no substantial task or group differences among real dyads overall or across sessions, and no substantial relationships between cardiac synchrony and any measure of behaviour. We found trends of substantial relationships for intergenerational dyads between cardiac synchrony and drawing collaboration at all but four lags, with the four non-trending lags showing the same numerical relationship. We propose that these findings, considered together, suggest that cardiac synchrony is to a limited extent sensitive to higher stakes, open-ended, collaboration. We discuss the implications of these findings below.

### 4.1 Relationships between behavioural and self-report measures

Intergenerational and same generation dyads exhibited some similarities in behavioural measures alongside marked differences. For both groups, greater cumulative feelings of social closeness within dyads co-occurred with greater ratings of collaboration of the dyads’ drawings. This finding is consistent with previous findings ^[57–59]^. Notably, however, same generation dyads who reported more similar feelings of social closeness (as opposed to intergenerational dyads) completed the jigsaw puzzle more quickly. This pattern was not present in intergenerational dyads. We propose that drawing and puzzle assembly, both collaborative tasks, differ in the clarity of the goal. The goal of puzzle assembly is clear to both dyad members (i.e., correctly assemble all pieces before time runs out) while the goal of drawing may be more open-ended, based on the amount and specificity of dyads’ verbal planning prior to drawing and the process of co-drawing without talking. For reasons relating to experimental control, most studies to date have focused collaborative tasks with clear end goals ^[11,23,27,60]^. The value of the large dataset and analyses that we present here lies in their ecological validity. That is, they reflect everyday collaboration arguably more accurately than some laboratory-based measures reported by previous studies. Everyday collaboration inherently entails free choice regarding the extent to which one engages and mutual adaptation to one’s partner, which have consequences for the development of social relationships.

We did not expect groups to differ in how close they sat and/or leaned toward one another while drawing together. Yet, intergenerational dyads drew sitting closer together than same generation dyads. Further, the relationships between interpersonal distance and feelings of social closeness, measured as a cumulative score per dyad and the discrepancy between dyad members, was opposing between groups. Intergenerational dyads sat closer when they felt more socially close or had smaller discrepancies in their feelings of social closeness–showing the pattern we anticipated based on existing findings ^[32–34]^. Same generation dyads, on the other hand, sat closer when they felt less socially close or had greater discrepancies in their feelings of social closeness. These opposite patterns suggest that reduced levels of social comfort during collaboration may amplify dyads’ perceptions of the stakes of the collaboration and result in compensatory behaviours, such as sitting closer to one another. Previous works corroborates this interpretation ^[61–64]^. Thus, interpersonal distance in collaboration may be a promising objective measure of social closeness and relationship development within recently acquainted dyads.

### 4.2 Cardiac synchrony not sensitive to true interaction, unless stakes are increased

Our analyses of cardiac synchrony revealed no substantial differences between real and pseudo dyads overall or across sessions, no substantial task or group differences among real dyads overall or across sessions, and no substantial relationships between cardiac synchrony and any measure of behaviour. This was true for averaged and separate lags. The absence of differences between real and pseudo dyads can be interpreted as meaning that any two individuals meeting the inclusion criteria of our study (whether same generation or intergenerational), who drew following the same prompts given by the researchers, in the same context are likely to exhibit comparable levels of cardiac synchrony, whether or not they are in the same room at the same time or acquainted. In other words, the levels of cardiac synchrony observed in this study do not index *true* interaction between two people. Moreover, the levels of cardiac synchrony observed in this study do not index changes in collaborative behaviour or feelings of social closeness across repeated encounters.

A small number of studies have previously reported that cardiac synchrony does not differ between real and pseudo dyads ^[60,65]^. Flory et al. ^[65]^ present alternative explanations. One is that cardiac synchrony is a spurious finding (i.e., could occur for any two people in the world). We do not think this interpretation applies here as the experimental setting, structure, and inclusion criteria provide substantial structure to the experience of each participant–that is, the description of cardiac synchrony as completely spurious is not founded. Another of Flory et al.’s alternative explanations is that participants operate in a common psychophysiological mode (i.e., similar mental state evoked by the experimental design), aligning with Danyluck and Page-Gould’s ^[11(p. 1)]^ view that “physiological synchrony could simply reflect the mutually experienced demands of a shared environment”. While this is in line with our rationale against spurious cardiac synchrony, we disagree on the basis that we observed trending relationships between cardiac synchrony and drawing collaboration for intergenerational dyads for 21 of 25 lags. We propose that these relationships point to a noteworthy, albeit non-substantial, influence of high stakes collaboration on cardiac synchrony.

We observed no task or group differences in cardiac synchrony measured within real dyads. Prior research has also reported the absence of task-specific or performance-related changes in cardiac synchrony ^[21,22,27,66]^. More specifically, Gordon et al. ^[22]^ detected evidence for task differences in cardiac synchrony when participants drummed together at a predetermined speed or drummed freely, improvising together. Similarly, other authors report having found no evidence to support associations between cardiac synchrony and task performance. Boukarras et al. ^[21]^ report that reaction times in a dyadic grasping task are not associated with cardiac synchrony. Behrens et al. ^[66]^ report that cooperative success is not associated with cardiac synchrony when dyads played the Prisoners Dilemma game. Strang et al. report that cooperative success is not associated with cardiac synchrony when dyads play Tetris with one person controlling object rotation and the other controlling the positioning ^[60]^. Finally, with respect to measures of social connection and closeness, heart rate coordination (consisting of alternate measures to cardiac synchrony) during Lego building was reported not to be associated with measures of perceived group relatedness or competence, but rather with speech and movement dynamics ^[27]^. Interested readers can find further examples in Mayo and Gordon’s review ^[26]^. In summary, despite the large number of studies dedicated to understanding cardiac synchrony on the grounds that cardiac synchrony can offer a window into collaboration and social connection, the extent to which this is true remains a topic worthy of debate (as well as replication).

From a practical perspective, the consistency of non-differences in cardiac synchrony across our preregistered exploratory analyses suggest that cardiac synchrony is not an optimal measure for predicting collaborative outcomes or relationship development in real-world environments. Given the mounting interest expressed in using wearable devices to collect objective insights relating to social wellbeing ^[67–70]^, we offer the following recommendation: Organisations, policy makers, and researchers aspiring to use wearable physiological monitoring to track changes in social behaviours and social wellbeing (i.e., social connection) should consider the low explanatory power of cardiac synchrony demonstrated here and pilot extensively before dedicating funding to large-scale roll-outs of these devices to measure program engagement or success. We feel justified in making this recommendation based on the sample size (366 samples involving within-participant/dyad design, which reduces inter-individual variability in our analyses). Moreover, other work by our research group shows that inter-brain synchrony may index changes in social wellbeing with greater sensitivity ^[8]^, though we acknowledge the logistical and financial barriers to larger-scale introduction of brain imaging measures.

Nonetheless, from an empirical perspective, it is promising that collaborative drawing scores show a trending relationship with cardiac synchrony levels within real intergenerational dyads. This suggests that further research could uncover which social, cognitive, and behavioural conditions need to be met for cardiac synchrony to be sensitive to true dyadic interaction. Based on the behavioural data reported here, we believe these conditions should include higher perceived stakes for successful collaboration. Our behavioural data revealed that intergenerational dyads’ drawings were rated more collaborative than same generation dyads’, and that the elevated levels of drawing collaboration emerged from the group (i.e., intergenerational dyads) that reported lower feelings of social closeness within dyads, more disparate feelings of social closeness within dyads, and sat closer together while drawing together (Table 1). In concert, these behaviours highlight how lower levels of social comfort during collaboration may raise the stakes of the collaboration and may, in turn, contribute to shared fluctuations in arousal during collaboration.

### 4.3 Limitations

In using the term “higher stakes collaboration”, we are attempting to articulate the origin of the decreased social comfort, or increased social tension, as it relates to reputation management ^[71,72]^ or the additional cognitive load stemming from interpersonal uncertainty ^[i.e., lack of common ground:, 73,74]^. We are open to this being conceptualised in other ways. Follow up experiments could probe the extent to which individual strategies and subjective experiences shed light on the nature of such collaboration.

Measures of dyadic synchrony can influence observed synchrony levels ^[75]^. We considered this carefully before preregistering our planned pipeline to calculate IBI time series. We compared an cardiac synchrony approach implemented previously with fNIRS ^[76]^ with more common IBI approaches ^[21,29]^. We opted for IBI to align our work with existing studies examining cardiac synchrony. Future research could use non-linear methods such as cross recurrence quantification analysis (CQRA), which is suited for non-stationary signals such as IBI ^[77]^. With respect to our comparisons of real and pseudo dyads’ levels of cardiac synchrony, a complementary approach would be to employ segment-shuffled surrogates within each dyad. In this method, each person’s cardiac time series is cut into short segments that are then randomly reordered so that the overall signal properties are preserved but true timing is disrupted ^[e.g., 15]^. This would reveal whether true dyads display cardiac synchrony on top of task-driven similarity. Duration of cardiac signals has been demonstrated to influence the likelihood of finding significant cardiac synchrony, particularly for smaller group sizes such as dyads ^[78]^. From this perspective, our design is optimised to identify true cardiac synchrony, should it be present. Nonetheless, future studies could use dedicated heart-rhythm monitoring, such as electrocardiograms, to examine free-choice, unstructured collaboration in intergenerational dyads. In suggesting this, we recognise that a complete replication of this study is likely not feasible, and we support smaller targeted replication efforts.

## 5 Conclusion

We charted cardiac synchrony, collaborative behaviour, social closeness and interpersonal distance within intergenerational and same generation dyads across six sessions involving a collaborative drawing task. Our analyses of collaborative behaviour, self-reported social closeness, and interpersonal distance, when considered together, suggest that social comfort drives relationships between social closeness, interpersonal distance and collaboration. Of these, interpersonal distance emerged as a promising objective measure of relationship development. We found that cardiac synchrony did not co-vary substantially with group, task, an interaction thereof or collaborative behaviour, social closeness or interpersonal distance. Nonetheless, we did find a trending relationship between collaboration while drawing together and cardiac synchrony that was specific to intergenerational dyads. These findings have implications for understanding the mechanisms underpinning social interaction within and between generations and for the implementation of cardiac synchrony as an objective measure of social connection or collaboration in everyday, real-world contexts.

## Supporting information

Supplementary Materials 26 Feb 2026

## Author contributions

RM: Conceptualisation, Equipment donation, Data curation, Formal analysis, Investigation, Visualisation, Writing – original draft, Writing – review & editing. LN: Data curation, Validation, Writing – original draft. ESC: Conceptualisation, Funding acquisition, Project administration, Writing – review & editing.

## Data availability statement

These data were collected after the submission of the following preregistration: https://osf.io/xuvy5. The data collected and analysed for this study can be found in our repository entitled ‘Cardiac synchrony during collaborative drawing: Intergenerational and longitudinal perspectives’ on OSF: https://osf.io/akmqw.

## Acknowledgements

We thank the survey center operated by UZH Healthy Longevity Center, University of Zurich, for their assistance with recruitment of older adults from the Zurich community. We thank Tessa Portier and Medea Häuselmann for their assistance with data collection, Linda Fanconi for scheduling participants’ sessions, Alistair Gadola for coding participant behaviour, and Fanny Mougel for digitising the drawings. We thank Chantal Nagel for assisting with literature review. We thank Courtney Casale and Tessa Portier for their help with proofreading.

