## Supplementary Materials 26 Feb 2026 for "Cardiac synchrony during collaborative drawing: A longitudinal comparison of same generation and intergenerational dyads"

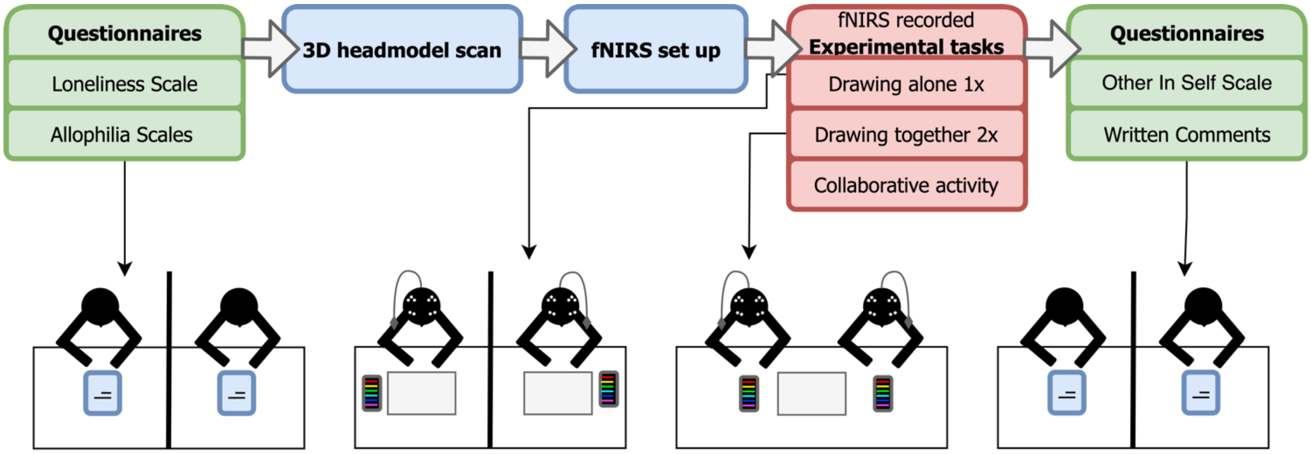


**Figure 1.** Flow chart showing the protocol employed at each session.

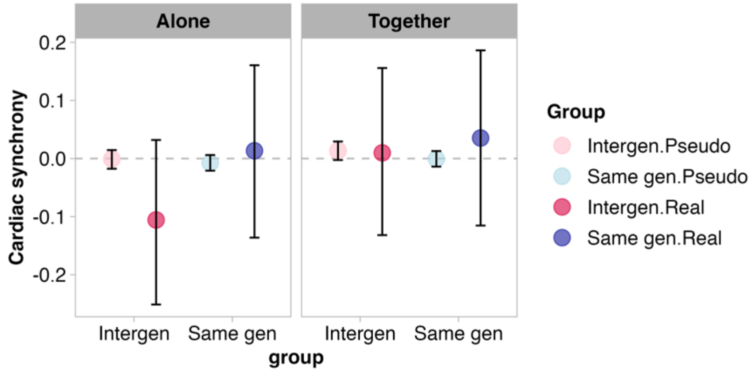


**Figure 2.** All sessions combined, parameter estimates per group and condition. Error bars show 95 % HPD.

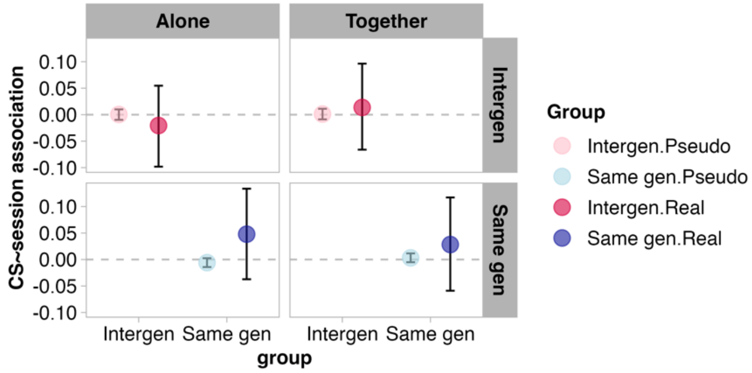


**Figure 3.** Change in cardiac synchrony (CS) across sessions, parameter estimates per group and condition. Error bars show 95 % HPD.

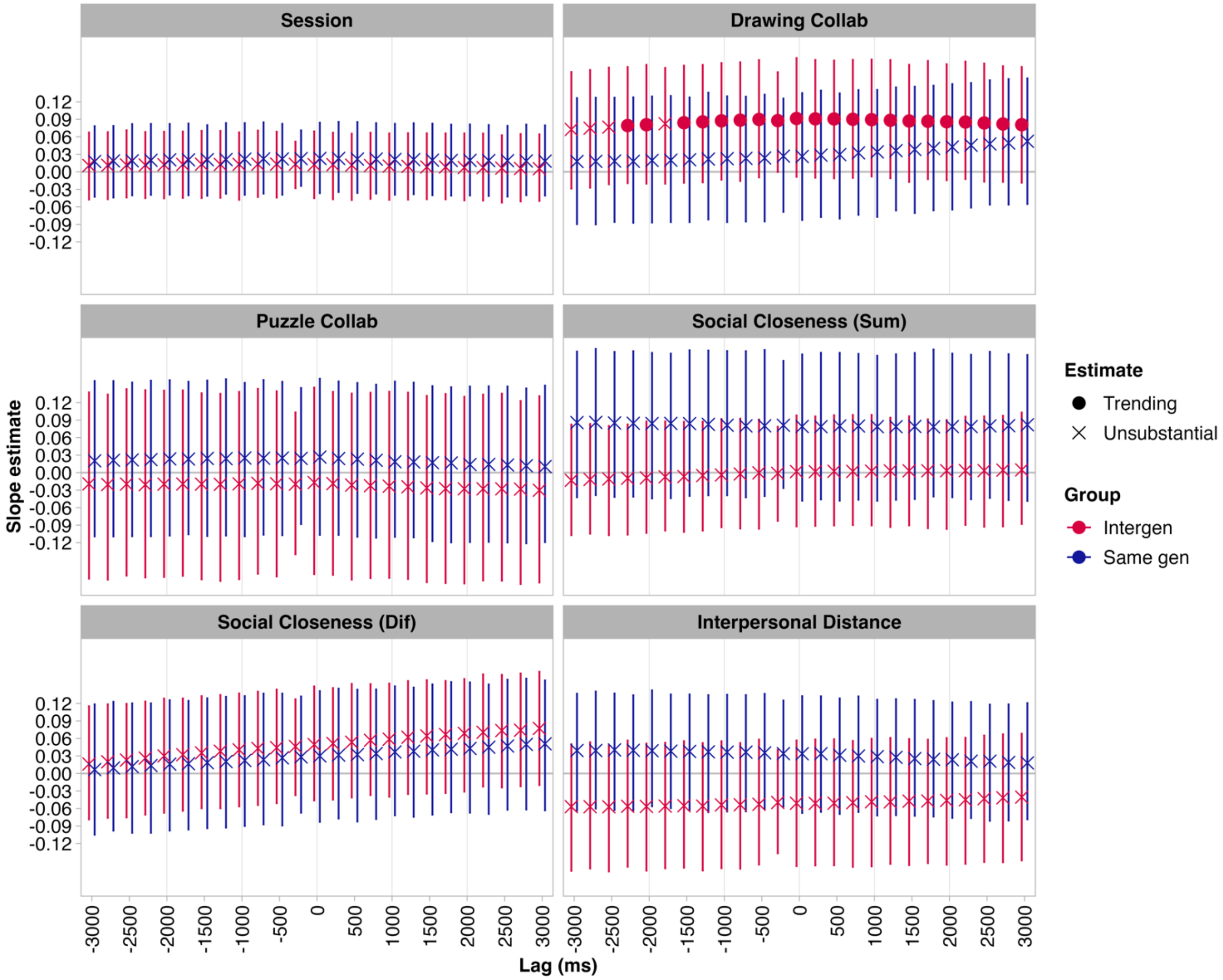


**Figure 4.** Relationship between cardiac synchrony and measures. Error bars show 95 % HPD.

**Table 1.** Breakdown of reasons for excluding dyads data. Recordings/dyads only included once in list. E.g., if a recording/dyad is counted among reasons unrelated to signal quality, they are not re-counted among reasons related to signal quality. In total 91 recordings of 732 recordings were excluded, impacting 85 of 366 dyadic sessions.

| **Reason** | **N individual recordings excluded** | **N dyadic sessions impacted** |
|---|---|---|
| ***Unrelated to signal quality*** | | |
| Recordings lost in transition between recording laptops | 2 | 2 |
| Triggers missing or unreliable | 7 | 4 |
| Participant unable to complete session (could not attend final session/consumed alcohol) | 5 | 3 |
| Multi-part recording (bluetooth or battery dropout) | 6 | 4 |
| ***Related to signal quality*** | | |
| No short channels of adequate quality | 71 | 69 |
| No long channels of adequate quality (pilot montage used) | 0 | 0 |
| **Total** | **91** | **85** |

**Table 2.** Standardised estimates of relationship between behavioural and self-report measure per group.

| | **Drawing Collaboration** | **Puzzle Collaboration** | **Social Closeness (Sum)** | **Social Closeness (Dif)** |
|---|---|---|---|---|
| **Drawing Collaboration (Intergen)** | - | | | |
| **Drawing Collaboration (Samegen)** | - | | | |
| **Puzzle Collaboration (Intergen)** | 0.19 [0.06, 0.32] | - | | |
| **Puzzle Collaboration (Samegen)** | -0.07 [-0.18, 0.04] | - | | |
| **Social Closeness Sum (Intergen)** | 0.19 [0.10, 0.28] | -0.05 [-0.13, 0.02] | - | |
| **Social Closeness Sum (Samegen)** | 0.35 [0.23, 0.46] | -0.23 [-0.33, 0.13] | - | |
| **Social Closeness Dif (Intergen)** | -0.03 [-0.13, 0.08] | 0.06 [-0.02, 0.14] | 0.19 [0.11, 0.27] | - |
| **Social Closeness Dif (Samegen)** | -0.00 [-0.10, 0.11] | -0.15 [-0.23, -0.06] | 0.11 [0.02, 0.19] | - |
| **Proximity (Intergen)** | -0.11 [-0.22, -0.01] | -0.02 [-0.11, 0.05] | 0.10 [0.01, 0.19] | -0.09 [-0.19, 0.01] |
| **Proximity (Samegen)** | 0.10 [0.00, 0.20] | -0.09 [-0.17, -0.00] | -0.13 [-0.24, -0.01] | 0.12 [0.02, 0.22] |
